# Phylogenetic confidence intervals for the optimal trait value

**DOI:** 10.1101/005314

**Authors:** Krzysztof Bartoszek, Serik Sagitov

## Abstract

We consider a stochastic evolutionary model for a phenotype developing amongst *n* related species with unknown phylogeny. The unknown tree is modelled by a Yule process conditioned on *n* contemporary nodes. The trait value is assumed to evolve along lineages as an Ornstein-Uhlenbeck process. As a result, the trait values of the *n* species form a sample with dependent observations. We establish three limit theorems for the sample mean corresponding to three domains for the adaptation rate. In the case of fast adaptation, we show that for large n the normalized sample mean is approximately normally distributed. Using these limit theorems, we develop novel confidence interval formulae for the optimal trait value.

## 1 Introduction

Phylogenetic comparative methods deal with multi-species trait value data. This is an established and rapidly expanding area of research concerning evolution of phenotypes in groups of related species living under various environmental conditions. An important feature of such data is the branching structure of evolution causing dependence among the observed trait values. For this reason the usual starting point for phylogenetic comparative studies is an inferred phylogeny describing the evolutionary relationships. The likelihood can be computed by assuming a model for trait evolution along the branches of this fixed tree, such as the ornstein-Uhlenbeck process.

The one-dimensional Ornstein-Uhlenbeck model is characterized by four parameters: the optimal value *θ*, the adaptation rate *α* > 0, the ancestral value *X*_0_, and the noise size *σ*. The classical Brownian motion model [14] can be viewed as a special case with *α* = 0 and *θ* being irrelevant. As with any statistical procedure, it is important to be able to compute confidence intervals for these parameters. However, confidence intervals are often not mentioned in phylogenetic comparative studies [8].

There are a number of possible numerical ways of calculating such confidence intervals when the underlying phylogenetic tree is known. Using a regression framework one can apply standard regression theory methods to compute confidence intervals for (*θ, X*_0_) conditionally on (*α*, *σ*^2^) [15, 20, 28, 33]. Notably in [16] the authors derive analytical formulae for confidence intervals for *X*_0_ under the Brownian motion model. In more complicated situations a parametric bootstrap is a (computationally very demanding) way out [8, 11, 27]. Another approach is to report a support surface [20, 21], or consider the curvature of the likelihood surface [7].

All of the above methods have in common that they assume that the phylogeny describing the evolutionary relationships is fully resolved. Possible errors in the topology can cause problems-the closer to the tips they occur, the more problematic they can be [43]. On the other hand, the regression estimators will remain unbiased even with a misspecified tree [34] and also seem to be robust with respect to errors in the phylogeny at least for the Brownian motion model [42]. There are only few papers addressing the issue of phylogenetic uncertainty. An MCMC procedure to jointly estimate the phylogeny and parameters of the Brownian model of trait evolution was suggested in [26, 25]. Recently, [36] develops an Approximate Bayesian Computation framework to estimate Brownian motion parameters in the case of an incomplete tree.

Our paper studies a situation when nothing is known about the phylogeny. The simplest stochastic model addressing this case is a combination of a Yule tree and the Brownian motion on top of it: already in the 1970s, a joint maximum likelihood estimation procedure of a Yule tree and Brownian motion on top of it was proposed in [13]. This basic evolutionary model allows for far reaching analytical analysis [6, 12, 35]. A more realistic stochastic model of this kind combines the Brownian motion with a birth-death tree allowing for extinction of species [10]. For the latter model [35] explicitly compute the so-called interspecies correlation coefficient. Such “tree-free” models are appropriate for working with fossil data when there may be available rich fossilized phenotypic information but the molecular material might have degraded so much that it is impossible to infer evolutionary relationships. In [12] the usefulness of the tree-free approach for contemporary species is demonstrated in an Carnivora order case study and in [31] the distribution over the space of Yule trees of the interspecies correlation coefficient is calculated.

Conditioned birth-death processes as stochastic models for species trees, have received significant attention in the last decade [3, 18, 29, 38, 39, 40]. In this work the unknown tree is modeled by the Yule process conditioned on *n* extant species while the evolution of a trait along a lineage is viewed as the Ornstein-Uhlenbeck process, see Fig. 1. We study the properties of the sample mean and sample variance computed from the vector of *n* trait values. Our main results are three asymptotic confidence interval formulae for the optimal trait value *θ*. These three formulae represent three asymptotic regimes for different values of the adaptation rate *α*.

In the discussion in [12] it is pointed out that “as evolutionary biologists further refine our knowledge of the tree of life, the number of clades whose phylogeny is truly unknown may diminish, along with interest in tree-free estimation methods.” In our opinion the main contribution of such methods is that they indicate statistical and asymptotic properties of phylogenetic samples under given evolutionary models. These properties can then be verified for other models of tree growth or real phylogenies [4, 5, 17, 22, 23, 24, 30]. We believe furthermore that the easy-to-compute tree-free predictions will always play an important role of a sanity check to see whether the conclusions based on the inferred phylogeny deviate much from those from a “typical” phylogeny. Moreover, results like those presented here can also be used as a method of testing software for phylogenetic comparative models.

A detailed description of the evolutionary model along with our main results are presented in Section 2. Section 3 contains new formulae for the Laplace transforms of important characteristics of the conditioned Yule species tree: the time to origin *U_n_* and the time *τ*^(*n*)^ to the most recent common ancestor for a pair of two species chosen at random out of *n* extant species. In Section 4 we calculate the interspecies correlation coefficient for the Yule-Ornstein-Uhlenbeck model and Section 5 contains the proof of our limit theorems. In Section 6 we establish the consistency of the stationary variance estimator, which is needed for our confidence interval formulae, cf [20] where the residual sum of squares was suggested to estimate the stationary variance. In Appendix A we calculate all the joint moments of *U_n_* and *τ*^(*n*)^.

Our main result, Theorem 2.1, should be compared with the limit theorems obtained in [1, 2]. They also revealed three asymptotic regimes in a related, though different setting, dealing with a branching Ornstein-Uhlenbeck process. In their case the time of observation is deterministic and the number of the tree tips is random, while in our case the observation time is random and the number of the tips is deterministic. Although it is possible (with some effort) to deduce our results from [1, 2], our proof provides a much more elementary derivation. We believe that our approach will be useful in addressing other biologically relevant issues like the formulae for the higher moments given in Appendix A. Another similar limit theorem, but one conditional on the sequence of species trees generated by different mechanisms, is derived in [5].

**Figure 1:**
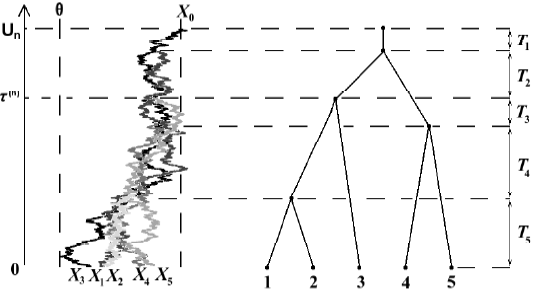
On the left: a branching Ornstein-Uhlenbeck process simulated on a realization of the Yule *n*-tree with *n* = 5 tips using the TreeSim [38, 39] and mvS-LOUCH [7] R [32] packages. Parameters used are *α* = 1, *σ* = 1, *X*_0_ −*θ* = 2, after the tree height *U_n_* was scaled to 1. On the right: the species tree disregarding the trait values supplied with the notation for the inter-speciation times. For the pair of tips (2,3) the time *τ*^(*n*)^ to their most recent common ancestor is marked on the time axis (starting at present and going back to the time of origin).

## 2 The model and main results

This work deals with what we call the Yule-Ornstein-Uhlenbeck model which is characterized by four parameters (*X_0_, α, σ, θ*) and consists of two ingredients

1. the species tree connecting *n* extant species is modeled by the pure birth Yule process [44] with a unit speciation rate λ = 1 and conditioned on having *n* tips [18],
2. the observed trait values 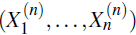 on the tips of the tree evolved from the ancestral state *X*_0_ following the Ornstein-Uhlenbeck process with parameters (*α*, *σ*, *θ*).

### Definition 2.1

*Let* (*T*_1_,…, *T*_*n*_) *be independent exponential random variables with parameters* (1,…, *n*). *We define the Yule n-tree as a random tree with n tips which is constructed using a bottom-up algorithm based on the following two simple rules*.

1. *During the time period *T*_*k*_ the tree has *k* branches*.
2. *For k ∈* [2, *n*] *the reduction from k to k* − 1 *branches occurs as two randomly chosen branches coalesce into one branch*.

*The height the Yule n-tree is now U_n_* = *T*_1_ + … + *T*_*n*_.

As shown in [18], this definition corresponds to the standard Yule tree conditioned on having *n* tips at the moment of observation, assuming that the time to the origin has the improper uniform prior.

Following [11, 20], we model trait evolution along a lineage using the Ornstein-Uhlenbeck process *X*(*t*) given by the stochastic differential equation

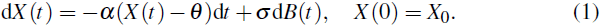

Here *α* > 0 is the adaptation rate, *θ* is the optimal trait value, *σ*^2^ is the noise variance, and *B*(*t*) is the standard Wiener process. The distribution of *X(t*) is normal with,

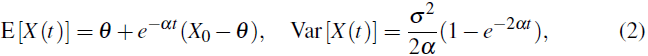

implying that *X*(*t*) looses the effect of the ancestral state *X*_0_ at an exponential rate. In the long run the Ornstein-Uhlenbeck process acquires a stationary normal distribution with mean *θ* and variance *σ*^2^/2*α*.

We propose asymptotic confidence interval formulae for the optimal value *θ* which take into account phylogenetic uncertainty. To this end we study properties of the sample mean and sample variance

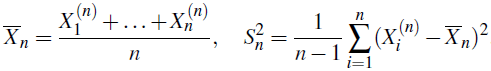

Using the properties of the Yule-Ornstein-Uhlenbeck model we find explicit expressions for 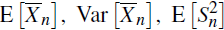, study the asymptotics of 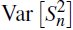, and prove the following limit theorem revealing three different asymptotic regimes.

### Theorem 2.1

*Let* 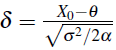 *be a normalized difference between the ancestral and optimal values. Consider the normalized sample mean 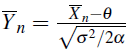 of the Yule-Ornstein-Uhlenbeck process with 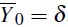. As *n* → ∞ the process 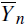 has the following limit behavior*.

i. *If α* > 0.5, *then 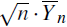 is asymptotically normally distributed with zero mean and variance* 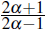.
ii. *If α* = 0.5, *then 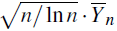 is asymptotically normally distributed with zero mean and variance* 2.
iii. *If α* < 0.5, *then 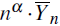 converges a.s. and in *L*^2^ to a random variable *Y*_α,δ_ with* 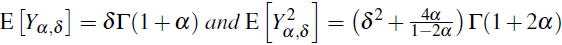.

Let *z_x_* be the *x*-quantile of the standard normal distribution, and *q_x_* be the *x*-quantile of the limit *Y_α,δ_*. Denote by *S_n_* the sample standard deviation defined as the square root of 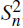. As it will be shown in Section 6, the sample variance 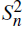 is a consistent estimator of 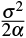. This fact together with Theorem 2.1 allows us to state the following three approximate (1 − *x*)-level confidence intervals for *θ* assuming that we know the value of *α*:

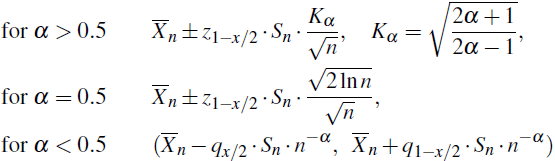

Notably, the first of these confidence intervals differs from the classical confidence interval for the mean 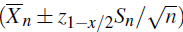 just by a factor *K_α_*. The latter is larger than 1, as it should, in view of a positive correlation among the sample observations. The correction factor *K_α_* becomes negligible in the case of a very strong adaptation, *α* ⪢ 1, when the dependence due to common ancestry can be neglected.

### Remark 2.1

*Observe that our standing assumption* λ = 1, *see Definition 2.1, of having one speciation event per unit of time causes no loss of generality. To incorporate an arbitrary speciation rate λ one has to replace in our formulae parameters α and σ*^2^ *by α/λ and σ*^2^/*λ. This transformation corresponds to the time scaling by factor λ in Eq. (1), it changes neither the optimal value θ nor the stationary variance σ*^2^/(2*α*).

## 3 Sampling *m* leaves from the Yule *n*-tree

Here we consider the Yule *n*-tree, see Definition 2.1 and study some properties of its subtree joining *m* randomly (without replacement) chosen tips, where *m* ∈ [2, *n*]. In particular, we compute the joint Laplace transform of the height of the Yule *n*-tree *U_n_* = *T*_1_ + … + *T_n_* and *τ*^(*n*)^, the height of the most recent common ancestor for two randomly sampled tips, see Fig. 1. For other results concerning the distribution of *τ*^(*n*)^ and *U_n_* see also [18, 19, 29, 35, 37, 38, 40, 41].

### Lemma 3.1

*Consider a random m-subtree of the conditioned Yule n-tree. It has m* − 1 *bifurcating events. Let 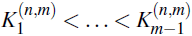 be the consecutive numbers of the bifurcation events in the Yule n-tree (counted from the root toward the leaves) corresponding to the m* − 1 *bifurcating events of the m-subtree. Put 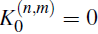 and 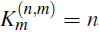. The sequence 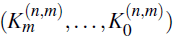 forms a time inhomogeneous Markov chain with transition probabilities*

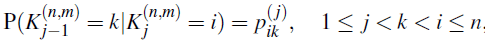

*where* 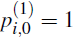 *for all i* ≥ 1, and

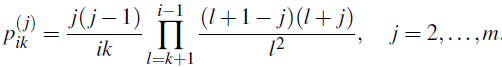

PROOF Tracing the lineages of *m* randomly sampled tips of the Yule *n*-tree towards the root, the first coalescent event can be viewed as the success in a sequence of independent Bernoulli trials. This argument leads to the expression cf [38]

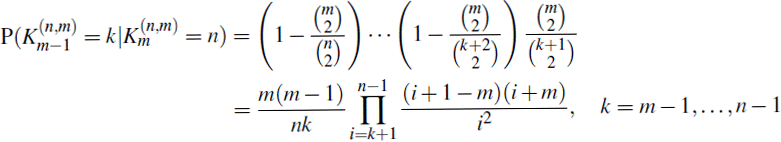

confirming the formula stated for the transition probabilities 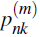. The transition probabilities 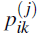 for *j* = 2,…, *m* − 1 are obtained similarly. Notice, as a check, that 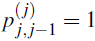.

### Lemma 3.2

*Consider the inter-bifurcation times for the m-subtree of the Yule n-tree*

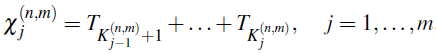

*so that 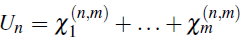 for any m* ≤ *n, and 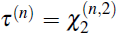 Then for x_j_* > − 1 *we have*

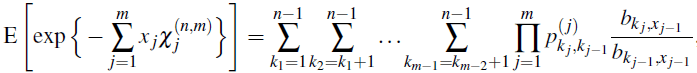

*where k_m_* = *n, k*_0_ = 0, *and*

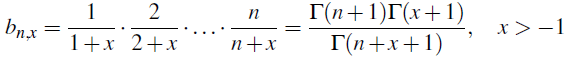

PROOF The Laplace transform of the sum of independent exponentials:

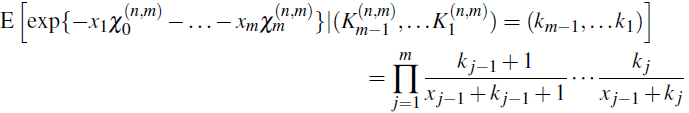

together with Lemma 3.1 implies the stated equality

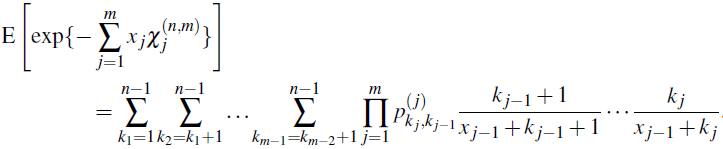

### Lemma 3.3

*The joint Laplace transform of the height of the Yule n-tree and the height of the most recent common ancestor for two randomly sampled tips is given by*

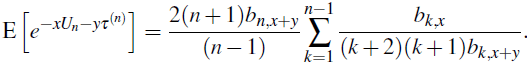

*In particular*,

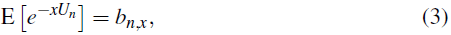

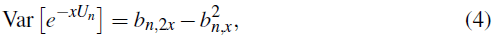

*and, denoting the harmonic number h_n_*: = 1 + 1/2 + … + 1/*n*,

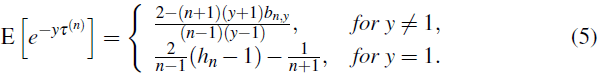

PROOF Turning to Lemma 3.1 with *m* = 2 we get

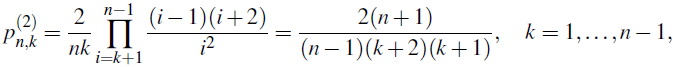

and according to Lemma 3.2

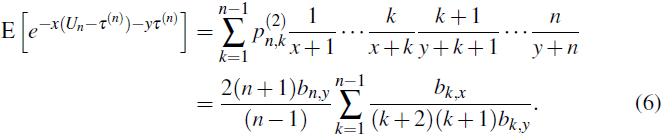

This implies the main formula claimed by Lemma 3.3 giving E[*e^−U_n_^*]= *b_n,x_* after putting *y* = 0. With *x* = 0,

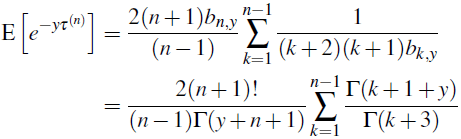

When *y* = 1 this directly becomes

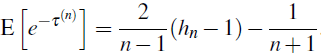

In the case of *y* ≠ 1 we use the following relation (easily verified by induction when *z* ≠ *y*)

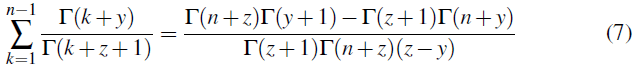

to derive

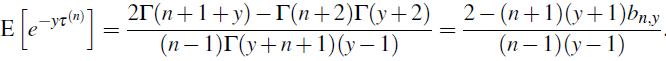

### Lemma 3.4

*As n* → ∞ *for positive x and y we have the following asymptotic results*

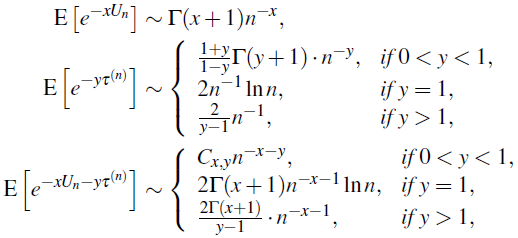

*where*

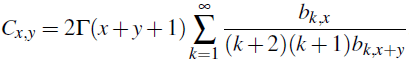

PROOF The stated results are obtained from Lemma 3.3 using the first of the following three asymptotic properties of the function *b_n,x_*

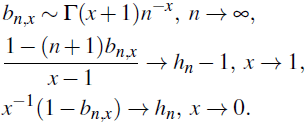

These three relations will often be used tacitly in what follows.

## 4 Interspecies correlation

Denote by 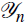 the *σ*-algebra containing all information on the Yule *n*-tree. The scaledtrait values 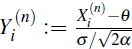 with in view of Eq. (2), are conditionally normal with

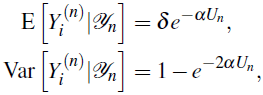

which together with the results from Section 3 entails

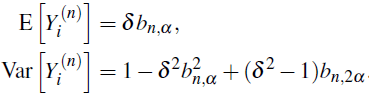

### Lemma 4.1

*In the framework of the Yule-Ornstein-Uhlenbeck model, for an arbitrary pair of traits we have*

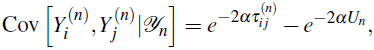

*where 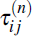 is the backward time to the most recent common ancestor of the tips (i, j)*.

PROOF Denote by 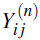 the normalized trait value of the most recent common ancestor of the tips (*i*, *j*). Let 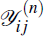 stand for the *σ*-algebra generated by 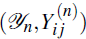, then using Eq. (2) we get

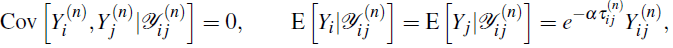

implying the statement of this lemma

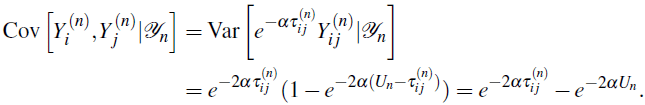

### Lemma 4.2

*Consider the interspecies correlation coefficient, the unconditioned correlation between two randomly sampled trait values*

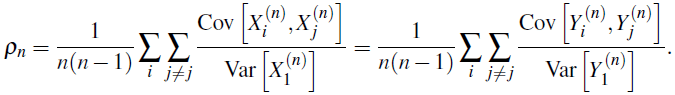

*If α* ≠ 0.5, *then*

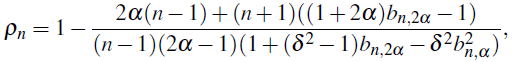

*and in the case of α* = 0.5

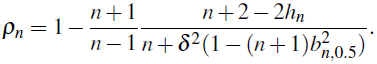

PROOF According to Lemma 4.1 we have,

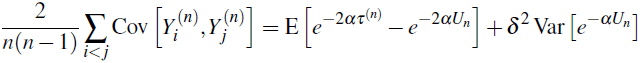

leading to

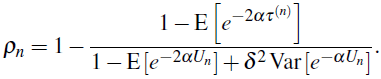

Applying the results of Section 3 we arrive at the asserted relations for *ρ_n_*. Observe that asymptotically as *n* → ∞ the interspecies correlation coefficient decays to 0 as

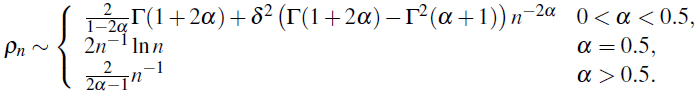

### Lemma 4.3

*Consider the sample mean 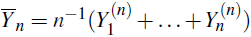 and the sample variance*

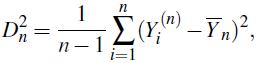

*of the scaled trait values. For all α* > 0 *we have* 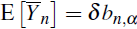. *For α* ≠ 0.5

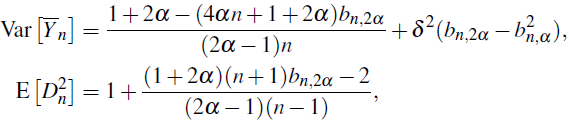

*and in the singular case α* = 0.5

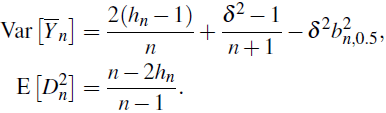

PROOF Obviously, 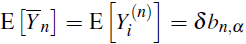. To prove the other assertions we turn to [35], where the concept of interspecies correlation was originally introduced. It was shown there that the variance of the sample average and the expectation of the sample variance can be compactly expressed as

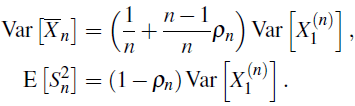

Since 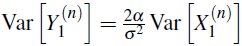, 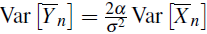, and 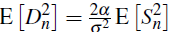 it remains to combine Lemma 4.2 with the known expression for 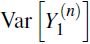.

A more direct proof of Lemma 4.3 can be obtained using the following result on conditional expectations.

### Lemma 4.4

*We have*

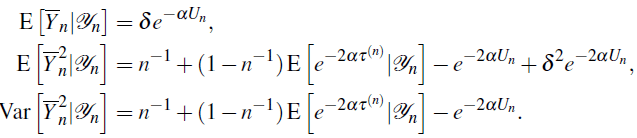

PROOF The main assertion follows from

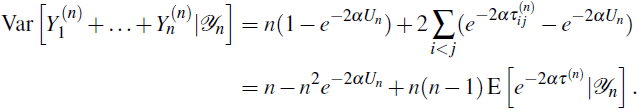

## 5 Proof of Theorem 2.1

### Lemma 5.1

*Put 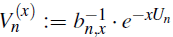 with 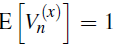. For any x* > −1 *the sequence 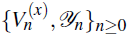 forms a martingale converging a.s. and in L*^2^. *Moreover*, (*U_n_* − log^*n*^) *converges in distribution to a random variable having the standard Gumbel distribution*.

PROOF The martingale property is obvious

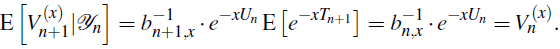

Since the second moments

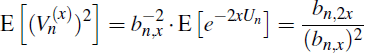

are uniformly bounded over *n*, we may conclude that 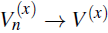 a.s. and in *L*^2^ with E[*V*^(*x*)^] = 1. It follows that 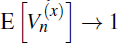, and therefore, E[*e*^−*x*(*U_n_* −log^*n*^)^] → Γ(*x* + 1). The latter is a convergence of Laplace transforms confirming the stated convergence in distribution.

Observe that the Gumbel limit for *U_n_* −log*n* can be obtained using the classical extreme value theory, in view of the representation

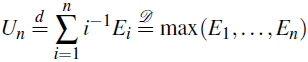

in terms of independent exponentials with parameter 1. Notice also that *U*_*n*+1_/2 has the same distribution as the total branch length of Kingman’s *n*-coalescent.

### Lemma 5.2

*Denote by ℱ_n_ the **σ**—algebra containing information on the Yule n-tree realization as well as the corresponding information on the evolution of trait values. Set*

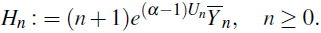

*The sequence 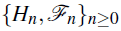 forms a martingale with* **E**[***H**_n_*] = *H*_0_ = *δ*.

PROOF Notice that,

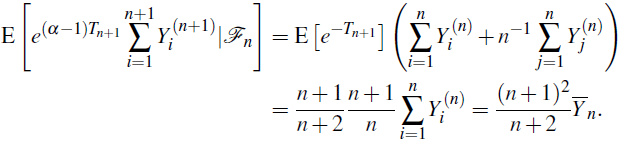

Hence

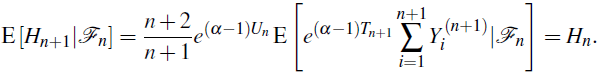

### Lemma 5.3

*For all positive α we have* 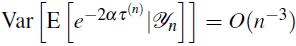 *as n* → ∞.

PROOF For a given realization of the Yule *n*-tree we denote by 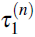 and 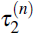 two independent versions of τ^(*n*)^ corresponding to two independent choices of pairs of tips out of *n* available. We have,

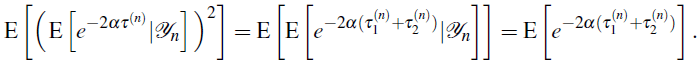

Writing

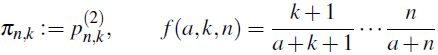

and using the ideas of Section 3 we obtain

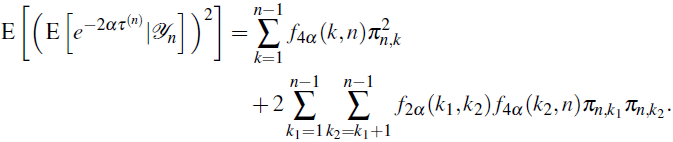

On the other hand,

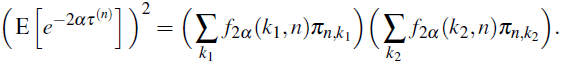

Taking the difference between the last two expressions we find

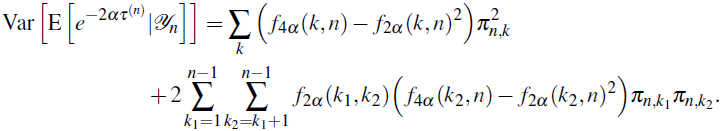

Using the simple equality

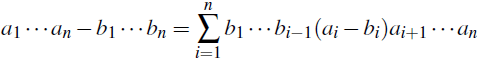

we see that it suffices to prove that,

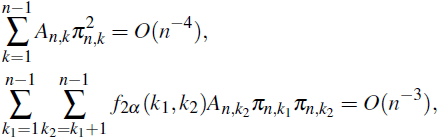

where

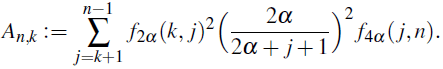

To verify these two asymptotic relations observe that

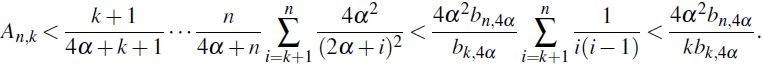

Since 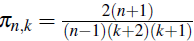, it follows

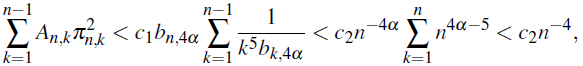

and

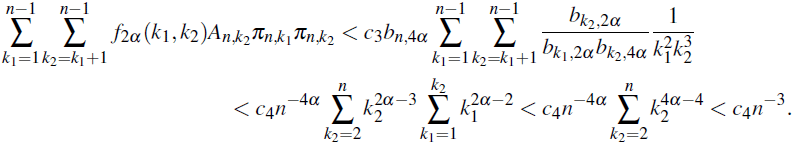

PROOF OF THEOREM 2.1 (I) AND (II). Let *α* > 0.5. To establish the stated normal approximation it is enough to prove the convergence in probability of the first two conditional moments

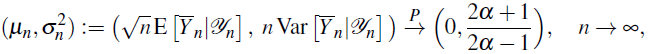

since then, due to the conditional normality of 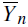, we will get the following convergence of characteristic functions

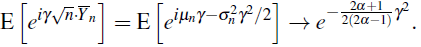

Now, due to Lemma 4.4 we can write

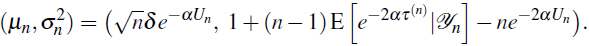

Using relations from Section 4 we see that

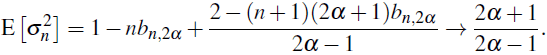

It remains to observe that on one hand, according to Lemma 5.3

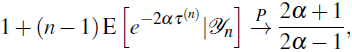

and on the other hand, 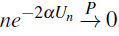 implying that 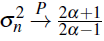. This together with *μ*_*n*_ → 0 holding in *L*^2^ and therefore in probability, entails 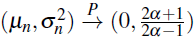, finishing the proof of part (i). Part (ii) is proven similarly.

PROOF OF THEOREM 2.1 (III). Let 0 < *α* < 0:5. Turning to Lemma 5.2 observe that the martingale 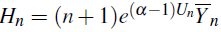 has uniformly bounded second moments. Indeed, due to Lemma 4.4

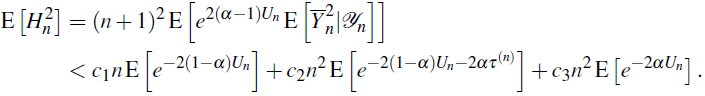

Thus, according to Lemma 3.4 we have 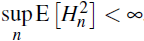. Referring to the martingale *L*^2^-convergence theorem we conclude that *H_n_* → *H_∞_* almost surely and in *L*^2^. Due to Lemma 5.1 it follows that

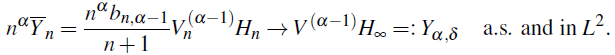

Finally, as *n* → ∞

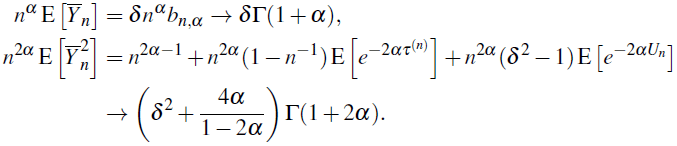

## 6 Consistency of the Sample Variance

Recall that 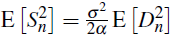, and according to Lemma 4.3 we have 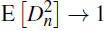. The aim of this section is to show that 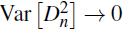 as *n* → ∞ which is equivalent to

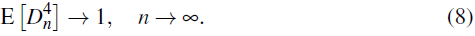

To this end we will need the following formula, see Eq. (13) in [9] valid for any normally distributed vector (Z_1_,Z_2_,Z_3_,Z_4_) with means (*m*_1_,*m*_2_,*m*_3_,*m*_4_) and covariances Cov [Z_*i*_, Z_*j*_] = *c_ij_*:

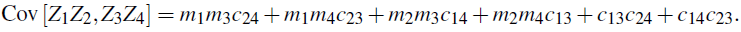

In the special case with *m_i_* = *m* it follows

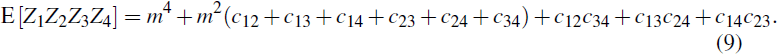

Writing *Y*_*i*_ instead of 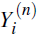 we use the representation

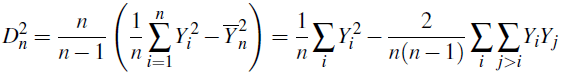

to find out that

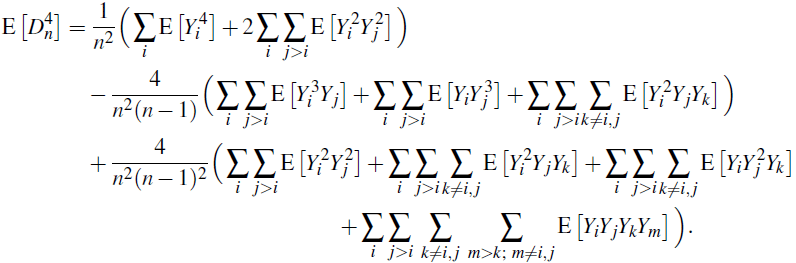

Denoting by (*W*_1_, *W*_2_, *W*_3_, *W*_4_) a random sample without replacement of four trait values out of *n* available, so that

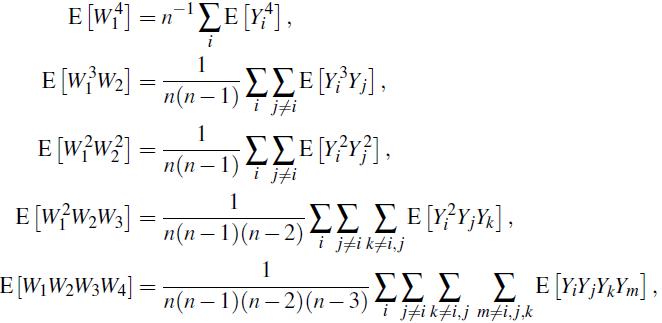

we derive

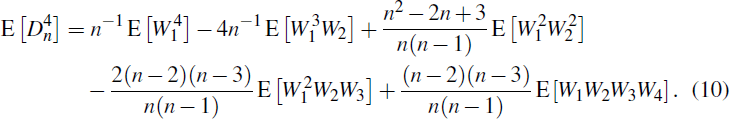

We compute the five fourth-order moments in the last expression using the conditional normality of the random quadruple (*W*_1_, *W*_2_, *W*_3_, *W*_4_) with conditional moments given by

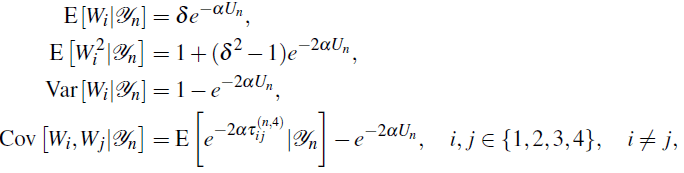

where 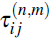 is the time to the most recent ancestor for the pair of tips (*i*, *j*) among *m* randomly chosen tips of the Yule *n*-tree. Clearly, all 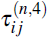 have the same distribution as τ^(*n*)^, and for

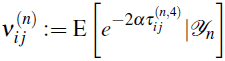

we can find the asymptotics of

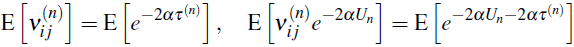

using Lemma 3.4. Notice also that

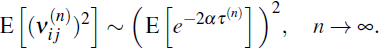

This follows from Lemma 5.3 and Lemma 3.4 as

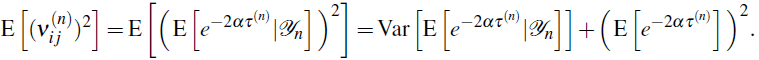

i. With Z_1_ = Z_2_ = Z_3_ = Z_4_ = *W*_1_ in Eq. (9), we obtain

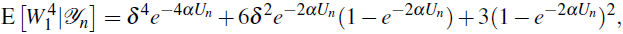

and therefore 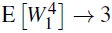 as n → ∞.
ii. Using Eq. (9) with Z_1_ = Z_2_ = Z_3_ = Z_4_ = *W*_1_ and Z_4_ = *W*_2_ we obtain

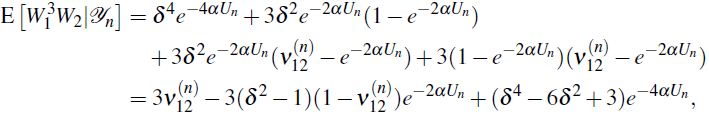

resulting in 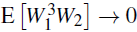 as *n* → ∞.
iii. Using Eq. (9) with Z_1_ = Z_2_ = *W*_1_ and Z_3_ = Z_4_ = *W*_2_ gives

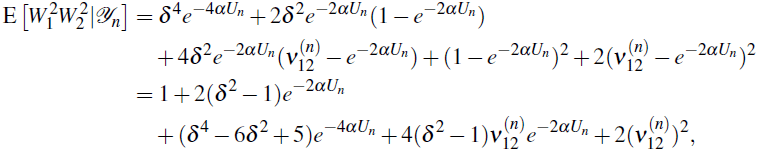

so that 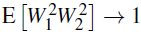 as *n* → ∞.
iv. Using a consequence of Eq. (9),

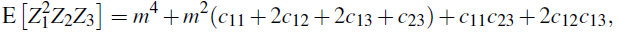

we get

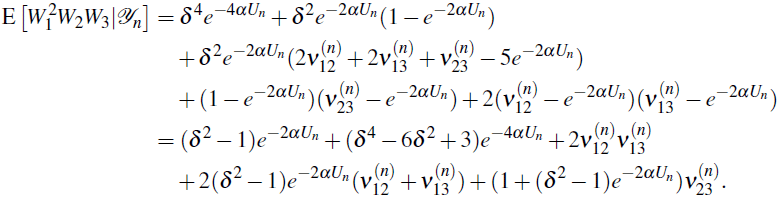 Using the Cauchy-Schwarz inequality

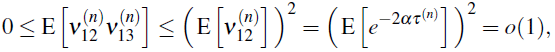

we obtain 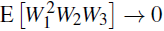 as *n* → ∞.
v. According to Eq. (9) we have

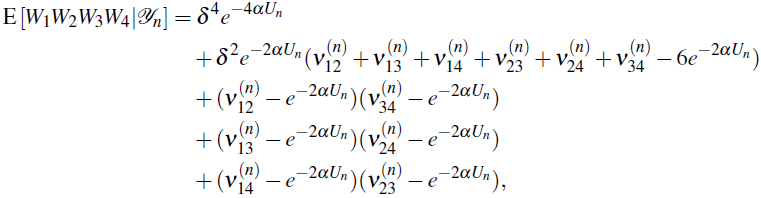

implying

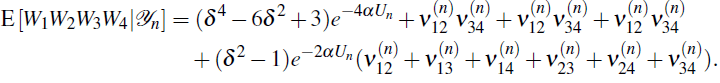 Using an estimate for 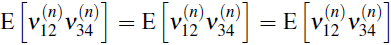 similar to that we used in (iv), we find E [*W*_1_*W*_2_*W*_3_*W*_4_] → 0 as *n* → ∞.

Finally, putting the above results (i)-(v) into Eq. (10) we arrive at Eq. (8).

## Acknowledgments

We are grateful to Thomas F. Hansen and Anna Stokowska for helpful suggestions and comments. Special thanks to an anonymous referee for the constructive suggestions to the earlier version of the paper, in particular, for the remark added after Lemma 5.1.

The research of Serik Sagitov was supported by the Swedish Research Council grant 621–2010–5623. Krzysztof Bartoszek was supported by the Centre for Theoretical Biology at the University of Gothenburg, Svenska Institutets Östersjösamarbetescholarship nr. 11142/2013, Stiftelsen för Vetenskaplig Forskning och Utbildning i Matematik (Foundation for Scientific Research and Education in Mathematics), Knut and Alice Wallenbergs travel fund, Paul and Marie Berghaus fund, the Royal Swedish Academy of Sciences, and Wilhelm and Martina Lundgrens research fund.

## A All Moments of *U_n_* and τ^(*n*)^

Eq. (3) for the Laplace transforms of the random variable *U_n_* can be used to calculate the moments of *U_n_* using,

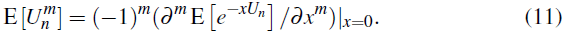

For a fixed *n* we introduce the following notation,

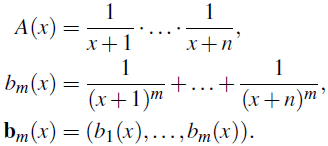

Notice that *A*(0) = 1/*n*! and *b_m_*(0) = *H*_*n*,*m*_ is the *n*-th generalized harmonic number of order *m*,

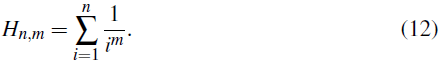

We can write Eq. (3) as E[*e*^−*xU_n_*^] = *n*!*A*(*x*). Its first derivative with respect to *x* is −*n*!*A*(*x*)*b*_1_(*x*), and the second derivative is *n*!*A*(*x*)(*b*_1_(*x*)^2^ + *b*_2_(*x*)). For the general recursive formula we introduce the following notation. We will denote by **k** = (*k*_1_, *k*_2_,…) infinite dimensional vectors with integer-valued components, and write 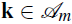 if all *k_i_* ≥ 0 and 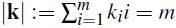. Therefore 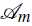 represents the set of all possible ways to represent *m* as a sum of positive integers. We will also use the multi-index notation **b**_*m*_(*x*)^**k**^ = *b*_1_ (*x*)^*k*_1_^ · … · *b_m_*(*x*)^*k_m_*^.

Since *A*′(*x*) = −*A*(*x*)*b*_1_(*x*), and *b*′*_m_*(*x*) = −*mb*_*m*+1_(*x*), we can show by induction that,

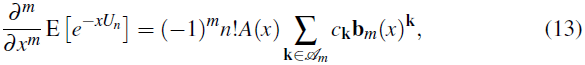

where coefficients *c*_**k**_ are defined for all vectors **k** = (*k*_1_,*k*_2_,…) with integer valued components using the recursion,

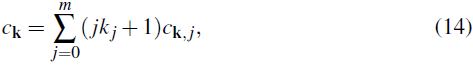

with *m* = |*k*| and

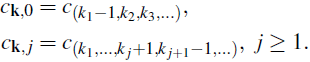

The boundary conditions for the recursion of Eq. (14) consist of two parts:

- *c*_**k**_ = 0, if all *k_i_* = 0, or one of the coordinates of the vector **k** is negative,
- *c*_**k**_ = 1 if *k*_1_ ≥ 1 and all other *k_i_* = 0.

We conclude from Eq. (13) that,

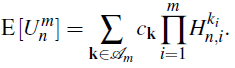

The technique for calculating the *m*-th derivative of the Laplace transform of τ^(*n*)^ given by Eq. (5) is the same but requires new notation

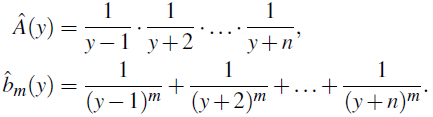

Notice that 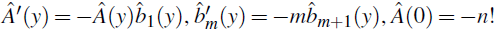 and 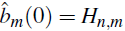 if *m* is even or 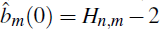 if *m* is odd. One can then inductively show that,

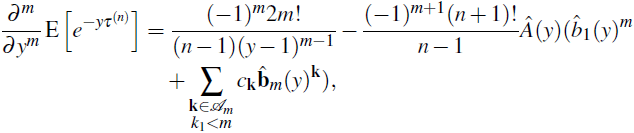

with the coefficients *c*_**k**_ defined as previously by Eq. (14). Therefore, we get,

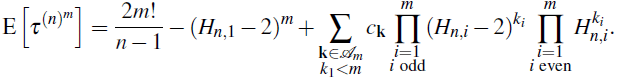

Similarly we can use Eq. (6) to calculate the joint moments for *U_n_* − τ^(*n*)^ and τ^(*n*)^ in terms of,

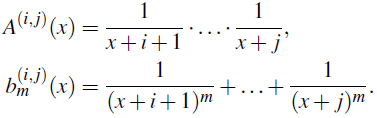

For *m* ≥ 1 and *r* ≥ 1 we first get,

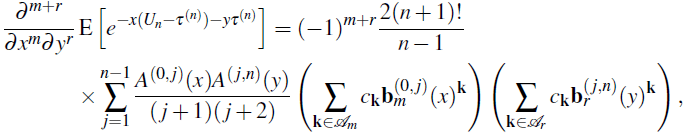

and then from the above,

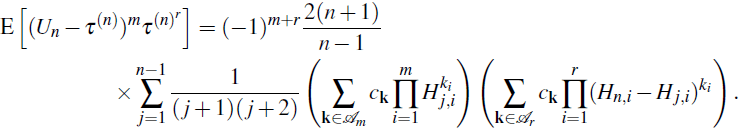

## References

[1] R. Adamczak and P. Milos. CLT for Ornstein-Uhlenbeck branching particle system. ArXiv e-prints, 2011.

[2] R. Adamczak and P. Milos. U-statistics of Ornstein-Uhlenbeck branching particle system. J. Th. Pobab., in press.

[3] D. Aldous and L. Popovic. A critical branching process model for biodiversity. Adv. Appl. Probab., 37(4):1094–1115, 2005.

[4] C. Ané. Analysis of comparative data with hierarchical autocorrelation. Ann. Appl. Stat, 2(3):1078–1102, 2008.

[5] C. Ané, L. S. T. Ho, and S. Roch. Phase transition on the convergence rate of parameter estimation under an Ornstein-Uhlenbeck diffusion on a tree. ArXiv e-prints, 2014.

[6] K. Bartoszek. Quantifying the effects of anagenetic and cladogenetic evolution. Math. Biosc., 254:42–57, 2014.

[7] K. Bartoszek, J. Pienaar, P. Mostad, S. Andersson, and T. F. Hansen. A phylogenetic comparative method for studying multivariate adaptation. J. Theor. Biol., 314:204–215, 2012.

[8] C. Boettiger, G. Coop, and P. Ralph. Is your phylogeny informative? Measuring the power of comparative methods. Evolution, 2012.

[9] G. W. Bohrnstedt and A. S. Goldberger. On the exact covariance of products of random variables. J. Am. Stat. Assoc., 64:1439–1442, 1969.

[10] F. Bokma. Time, species and seperating their effects on trait variance in clades. Syst. Biol., 59(5):602–607, 2010.

[11] M. A. Butler and A. A. King. Phylogenetic comparative analysis: a modelling approach for adaptive evolution. Am. Nat., 164(6):683–695, 2004.

[12] F. W. Crawford and M. A. Suchard. Diversity, disparity, and evolutionary rate estimation for unresolved Yule trees. Syst. Biol., 62(3):439–455, 2013.

[13] A. W. F. Edwards. Estimation of the branch points of a branching diffusion process. J. Roy. Stat. Soc. B, 32(2):155–174, 1970.

[14] J. Felsenstein. Phylogenies and the comparative method. Am. Nat., 125(1): 1–15, 1985.

[15] T. Garland and A. R. Ives. Using the past to predict the present: Confidence intervals for regression equations in phylogenetic comparative methods. Am. Nat., 155(3):346–364, 2000.

[16] T. Garland, P. E. Midford, and A. R. Ives. An introduction to phylogenetically based statistical methods, with a new method for confidence intervals on ancestral values. Amer. Zool., 39:374–388, 1999.

[17] O. Gascuel and M. Steel. Predicting the ancestral character changes in a tree is typically easier than predicting the root state. Syst. Biol., 63(3):421–435, 2014.

[18] T. Gernhard. The conditioned reconstructed process. J. Theor. Biol., 253: 769–778, 2008.

[19] T. Gernhard. New analytic results for speciation times in neutral models. B. Math. Biol., 70:1082–1097, 2008.

[20] T. F. Hansen. Stabilizing selection and the comparative analysis of adaptation. Evolution, 51(5):1341–1351, 1997.

[21] T. F. Hansen, J. Pienaar, and S. H. Orzack. A comparative method for studying adaptation to a randomly evolving environment. Evolution, 62:19651977, 2008.

[22] L. S. T. Ho and C. Ané. Asymptotic theory with hierarchical autocorrelations: Ornstein-Uhlenbeck tree models. Ann. Stat., 41(2):957–981, 2013.

[23] L. S. T. Ho and C. Ané. A linear-time algorithm for Gaussian and non-Gaussian trait evolution models. Syst. Biol., 63(3):397–408, 2014.

[24] L. S. T. Ho and C. Ané. Intrinsic inference difficulties for trait evolution with Ornstein-Uhlenbeck models. Meth. Ecol. Evol., in press.

[25] J. P. Huelsenbeck and B. Rannala. Detecting correlation between characters in a comparative analysis with uncertain phylogeny. Evolution, 57(6):1237–1247, 2003.

[26] J.P. Huelsenbeck, B. Rannala, and J.P. Masly. Accommodating phylogenetic uncertainty in evolutionary studies. Science, 88:2349–2350, 2000.

[27] A. R. Ives, P. E. Midford, and T. Garland. Within-species variation and measurement error in phylogenetic comparative methods. Syst. Biol., 56(2): 252–270, 2007.

[28] E. P. Martins and T. F. Hansen. Phylogenies and the comparative method: a general approach to incorporating phylogenetic information into the analysis of interspecific data. Am. Nat., 149(4):1341–1351, 1997.

[29] A. Mooers, O. Gascuel, T. Stadler, H. Li, and M. Steel. Branch lengths on birth-death trees and the expected loss of phylogenetic diversity. Syst. Biol., 61(2):195–203, 2012.

[30] E. Mossel and M. Steel. Majority rule has transition ratio 4 on Yule trees under a 2-state symmetric model. J. Theor. Biol., 360:315–318, 2014.

[31] W. H. Mulder and F. W. Crawford. On the distribution of interspecies correlation for Markov models of character evolution on Yule trees. J. Theor. Biol., 364:275–283, 2015.

[32] R Core Team. R: A Language and Environment for Statistical Computing. R Foundation for Statistical Computing, Vienna, Austria, 2013. URL http://www.R-project.org.

[33] F. J. Rohlf. Comparative methods for the analysis of continuous variables: geometric interpretations. Evolution, 55(11):2143–2160, 2001.

[34] F. J. Rohlf. A comment on phylogenetic correction. Evolution, 60(7):1509–1515,2006.

[35] S. Sagitov and K. Bartoszek. Interspecies correlation for neutrally evolving traits. J. Theor. Biol., 309:11–19, 2012.

[36] G. J. Slater, L. J. Harmon, D. Wegmann,P. Joyce, L. J. Revell, and M. E. Alfaro. Fitting models of continuous trait evolution to incompletely sampled comparative data using Approximate Bayesian Computation. Evolution, 66(1): 752–762, 2012.

[37] T. Stadler. Lineages-through-time plots of neutral models for speciation. Math. Biosci., 216:163–171, 2008.

[38] T. Stadler. On incomplete sampling under birth-death models and connections to the sampling-based coalescent. J. Theor. Biol., 261(1):58–68, 2009.

[39] T. Stadler. Simulating trees with a fixed number of extant species. Syst. Biol., 60(5):676–684, 2011.

[40] T. Stadler and M. Steel. Distribution of branch lengths and phylogenetic diversity under homogeneous speciation models. J. Theor. Biol., 297:33–40, 2012.

[41] M. Steel and A. McKenzie. Properties of phylogenetic trees generated by Yule-type speciation models. Math. Biosci., 170:91–112, 2001.

[42] E. A. Stone. Why the phylogenetic regression appears robust to tree mis-specification. Syst. Biol., 60(3):245–260, 2011.

[43] M. R. E. Symonds. The effects of topological inaccuracy in evolutionary trees on the phylogenetic comparative method of independent contrasts. Syst. Biol., 51(4):541–553, 2002.

[44] G. U. Yule. A mathematical theory of evolution: based on the conclusions of Dr. J. C. Willis. Philos. T Roy. Soc. B, 213:21–87, 1924.

